# Metabolite composition of sinking particles reflects a changing microbial community and differential metabolite degradation

**DOI:** 10.1101/459461

**Authors:** Winifred M. Johnson, Krista Longnecker, Melissa C. Kido Soule, William A. Arnold, Maya P. Bhatia, Steven J. Hallam, Benjamin A. S. Van Mooy, Elizabeth B. Kujawinski

## Abstract

Marine sinking particles transport carbon from the surface and bury it in deep sea sediments where it can be sequestered on geologic time scales. The combination of the surface ocean food web that produces these particles and the particle-associated microbial community that degrades these particles, creates a complex set of variables that control organic matter cycling. We use targeted metabolomics to characterize a suite of small biomolecules, or metabolites, in sinking particles and compare their metabolite composition to that of the suspended particles in the euphotic zone from which they are likely derived. These samples were collected in the South Atlantic subtropical gyre, as well as in the equatorial Atlantic region and the Amazon River plume. The composition of targeted metabolites in the sinking particles was relatively similar throughout the transect, despite the distinct oceanic regions in which they were generated. Metabolites possibly derived from the degradation of nucleic acids and lipids, such as xanthine and glycine betaine, were an increased mole fraction of the targeted metabolites in the sinking particles relative to surface suspended particles, while algal-derived metabolites like the osmolyte dimethylsulfoniopropionate were a smaller fraction of the observed metabolites on the sinking particles. These compositional changes are shaped both by the removal of metabolites associated with detritus delivered from the surface ocean and by production of metabolites by the sinking particle-associated microbial communities. Further, they provide a basis for examining the types and quantities of metabolites that may be delivered to the deep sea by sinking particles.

## Introduction

Sinking particles in the ocean are composed of a complex and varying mixture of dead or dying phytoplankton, zooplankton fecal pellets, aggregates held together by exopolysaccharides, and the microbes degrading this organic matter (Alldredge and Gotschalk 1989; Azam and Long 2001; Jackson 2001; Kiørboe and Jackson 2001; Roman et al. 2002; Durkin et al. 2016). These particles are responsible for delivering organic and inorganic carbon to deep ocean sediments where the carbon is buried on geologic time scales. Estimates from satellite data and models place the total global flux of carbon out of the euphotic zone at around 6 Pg C yr^-1^ (Siegel et al. 2014). However, the variables that control the magnitude of this flux in different ocean regions are still not fully understood. An important variable is the source and composition of particle-associated organic matter, and the mechanisms by which it is transformed and remineralized by particle-associated microbial communities (reviewed by Boyd and Trull 2007).

Marine particles are known hotspots of microbial activity. Microbes actively colonize these particles to exploit the abundance of bioavailable organic molecules (Kiørboe et al. 2002), resulting in a microbial community composition that is distinct from free-living communities (DeLong et al. 1993) and that expands the niche space for anaerobic metabolisms in the water column (Bianchi et al. 2018). The community that colonizes these particles consists of copiotrophic bacteria, specifically members of *Alteromonas, Methylophaga, Flavobacteriales*, and *Pseudoalteromonas*, known to be associated with dissolved organic matter enrichment or with eukaryotic phytoplankton (Fontanez et al. 2015). These microbes are well-adapted to utilize a diverse array of organic molecules (Fontanez et al. 2015) and thus exert an important control on the delivery of organic matter to the deep ocean.

During degradation of sinking particles, the relative contributions of major molecular classes of organic molecules do not change significantly with depth, while the total amount of organic carbon decreases (Hedges et al. 2001). Characterization of particulate organic matter (POM) at the level of molecular classes has shown that amino acids comprise approximately 40-50% of the major biomolecule classes contribution to total organic carbon, while 30% is carbohydrate and 20% is lipid (Hedges et al. 2001). However, the percentage of organic carbon that cannot be classified in a major class of biomolecules increases with depth, reaching ~80% in sea floor sediments (Wakeham et al. 1997b). Within classes of biomolecules, specific molecules are differentially degraded and produced on sinking particles. For instance, polar membrane lipids are degraded rapidly while other lipid types such as long-chain alkenones persist and reach the sediments (Wakeham et al. 1997a; Fulton et al. 2017). Similarly, some free amino acids like threonine, arginine, aspartic acid, glycine, and valine are enriched in deeper, more degraded sinking particle samples, suggesting that they are either not degraded as rapidly, or are produced as the particles sink (Ingalls et al. 2003). In contrast, relative monosaccharide composition on sinking particles is similar to the source composition with xylose and fucose decreasing slightly and arabinose increasing as a mole percent of carbohydrates (Tanoue and Handa 1980). While these studies come from many ocean regions and depths, they collectively show that there is measurable variability in concentrations of specific molecules despite relatively small variations in molecular class distributions across sinking marine POM.

Here, we use targeted metabolomics to investigate the dynamics of small biological molecules within sinking particles; specifically, we seek to understand how cellular components and metabolic products are cycled as particles exit the euphotic zone. Metabolomics is particularly well-suited to address this question, because this approach simultaneously characterizes a broad suite of small, structurally diverse organic molecules produced by biosynthetic pathways. These molecules include amino acids, nucleic acids, vitamins and cofactors, and biosynthetic intermediates but do not include biological macromolecules such as lipids, proteins, polysaccharides, or DNA/RNA. Using this approach, we can identify metabolic pathways or processes active during particle descent and degradation.

We compare the metabolite composition of sinking particles collected at 150 m in the western South and Equatorial Atlantic Ocean to that of suspended particles (i.e., cells) in the euphotic zone. Through this comparison of sinking and suspended particles (presumptive source for the sinking particles), we can characterize whether specific metabolites are removed during the course of particle production and descent or are produced in situ by particle-associated microbial communities. We hypothesize that as particles sink out of the euphotic zone, phytoplankton-derived metabolites are removed through microbial respiration and biochemical transformation, while metabolites associated with bacterial and archaeal biomass as well as those derived from organic matter degradation are added to the pool. This dataset identifies some metabolic indicators for these processes informing future field and laboratory studies.

## Materials and methods

### Field Sampling

We collected samples during two cruises: (1) KN210-04 in the western South Atlantic from 25 March – 9 May 2013, transiting from Montevideo, Uruguay to Bridgetown, Barbados and (2) AE1409 from Bermuda to Barbados between 8 May – 29 May 2014. We sampled suspended particles from water at three depths at each station: 5 m (referred to as the surface), the deep chlorophyll maximum (DCM; determined by fluorescence), and 250 m. To collect sinking particles, we used a conical net trap that was 1.25 m in diameter with a closed cod end (Peterson et al. 2005), deployed at 150 m and tethered to the surface. The trap was acoustically closed and retrieved 8–26 h after deployment (see Durkin et al. (2016) and Table S1 for deployment times). We split each net-trap sample into eight equal parts using a sample splitter (Lamborg et al. 2008), reserving one split for the metabolomics analysis and one for 16S rRNA sequencing.

### Materials

We purchased all metabolite standards from Sigma-Aldrich at the highest purity available with the following exceptions: dimethylsulfoniopropionate (DMSP), from Research Plus, Inc.; 2,3-dihydroxypropane-1-sulfonate (DHPS) and acetyltaurine, donated by Dr. Mary Ann Moran (University of Georgia); and *S*-(1,2-dicarboxyethyl)glutathione, from Bachem. We obtained hydrochloric acid (trace metal grade), acetonitrile (Optima grade), and methanol (Optima grade), from Fisher Scientific and formic acid from Fluka Analytical. We purchased glutamic acid-d_3_ from Cambridge Isotopes, 4-hydroxybenzoic acid-d4 from CDN Isotopes, and sodium taurocholate-d_5_ from Toronto Research Chemicals through Fisher Scientific. We purified all water with a Milli-Q system (Millipore; resistivity 18.2 MΩ·cm 25 °C, TOC < 1 *μ*M), and combusted all glassware used to collect samples and GF/F filters in an oven at 460 °C for at least 5 h before each cruise. Likewise, we autoclaved all plasticware, excluding bottles, tubing, and filter holders made of polytetrafluoroethylene (PTFE), before use. During the cruises, we flushed filter holders and tubing for shipboard filtration with 10% hydrochloric acid followed by Milli-Q water between each sample.

### Shipboard sample processing

We compared two types of particles (suspended particles and sinking particles) in this study that we defined operationally by their mode of collection. We sampled the suspended particles by collecting water samples (4 L) in a PTFE or polycarbonate bottles directly from Niskin bottles. We filtered each water sample sequentially through a GF/F filter (nominal pore size 0.7 *μ*m, Whatman) and a 0.2 *μ*m Omnipore filter (PTFE, Millipore) using a peristaltic pump. We collected sinking particles from the net trap splits in the same way, but filtering either through a GF/A filter (nominal pore size 1.6 *μ*m, stations K5, K7, K9, K11, K12, K18) or through a GF/F filter (stations K16, K19, K20, K21, K22, K23, A6, A12, A16) followed by a 0.2 *μ*m Omnipore filter (PTFE, Millipore). We stored all filters in their combusted aluminum foil wrappers at −80 °C until extraction in the laboratory.

### Laboratory sample processing

We extracted metabolites from filters as close to the time of analysis on the mass spectrometer as possible (< 48 h), using a protocol described by Rabinowitz and Kimball (2007) and modified for seawater samples (Kido Soule et al. 2015). One half of each filter from the suspended particle samples and 1/4 of the filter from the sinking particle samples was weighed and cut into small pieces using methanol-rinsed scissors and tweezers on combusted aluminum foil. Subsequently, we placed the filter pieces into an 8 mL glass amber vial with 1 mL −20 °C extraction solvent (40:40:20 acetonitrile:methanol:water + 0.1 M formic acid). Next, we added 25 *μ*L of a 1 *μ*g mL^−1^ deuterated standard mix (glutamic acid-d_3_, 4-hydroxybenzoic acid-d_4_, taurocholate-d_5_) as an extraction recovery standard. We gently vortexed each vial to separate filter pieces and expose them to the solvent; subsequently we sonicated the vials for 10 min. Then, we transferred the extracts to 1.5 mL Eppendorf centrifuge tubes with a Pasteur pipette. We rinsed the filter pieces in the 8 mL vials with 200 μL of cold extraction solvent and combined this rinse with the original transfer. We centrifuged the extracts at 20,000 x *g* for 5 min to remove cellular detritus and filter particles, and transferred the supernatant to clean 8 mL glass amber vials. We neutralized the extracts with 26 *μ*L of 6 M ammonium hydroxide and then dried the samples to near complete dryness in a Vacufuge (Eppendorf). Prior to mass spectrometry analysis, we reconstituted the suspended particle samples with either 247.5 *μ*L 95:5 water:acetonitrile and 2.5 *μ*L of 5 μg/mL biotin-d_4_ (injection standard) for samples collected at the surface or DCM or 123.8 *μ*L 95:5 water:acetonitrile and 1.25 *μ*L of 5 μg/mL biotin-d_4_ (injection standard) for deeper samples with presumed lower biomass. We reconstituted the sinking particle samples from the net trap in 495 *μ*L 95:5 water:acetonitrile and 5 *μ*L of 5 μg/mL biotin-d_4_ (injection standard). Finally, we transferred 100 *μ*L of each solution into individual glass inserts in autosampler vials for analysis on the mass spectrometer. For each sample type, we combined 15 *μ*L of each extract to create two pooled samples; one for suspended particle samples and one for sinking particle samples.

### Mass spectrometry

We performed HPLC-MS/MS analysis using a Phenomenex C18 column (Synergi Fusion, 2.1 × 150 mm, 4 *μ*m particle size) coupled via heated electrospray ionization (HESI) in polarity switching mode to a triple quadrupole mass spectrometer (Thermo Scientific TSQ Vantage) operated under selected reaction monitoring mode (SRM), as described previously (Kido Soule et al. 2015). We monitored quantification and confirmation ion SRM transitions for each analyte. We ran the following gradient at 250 *μ*L / min, with Eluent A (Milli-Q water with 0.1% formic acid) and Eluent B (acetonitrile with 0.1% formic acid): hold at 5% B for 2 min; ramp to 65% B for 16 min; ramp to 100% B for 7 minutes and hold for 8 min. We re-equilibrated the column with the starting ratio of eluents for 8.5 min between samples. We split the full run into batches of approximately 50 samples, and for each batch we used a pooled sample that corresponded to that batch. This allowed us to clean the instrument source and ion transfer tube at regular intervals, between batches. Before each batch, we conditioned the column with 5 injections of the pooled samples. The same pooled sample was then run after every 10 samples for quality control.

### Data processing

We converted XCalibur RAW files generated by the mass spectrometer to mzML files using MSConvert (Chambers et al. 2012), and we used MAVEN (Melamud et al. 2010; Clasquin et al. 2012) to select and integrate peaks. We discarded calibration peaks below a MAVEN quality threshold of 0.4 (on a scale of 0-1) and sample peaks below 0.2. These quality scores were generated by a machine-learning algorithm trained to recognize the quality of peaks by incorporating information from peaks scored by an expert human analyst (Melamud et al. 2010). To enhance confidence in metabolite identification, we required quantification and confirmation ion peaks to have retention times within 12 sec (0.2 min) of each other. We selected this range because peak width can range from 10–30 seconds and confirmation ions, which have lower intensity, can have a retention time assigned at a slightly different point on the peak resulting in retention times that can differ by ~0.1 minutes. Moreover, we required confirmation ions to have a MAVEN quality score of at least 0.1 and a signal-to-noise ratio greater than 1, and also that calibration curves have at least five calibration points with the highest calibration point one concentration level above the highest concentration in a sample. We adjusted concentrations based on ion suppression determined by comparison of a standard spiked pooled sample (1000 ng/mL metabolite standard mix in the sample matrix) to a standard spiked Milli-Q water sample (1000 ng/mL metabolite standard mix without complex matrix, see Table S2).

### Temperature, salinity, and chlorophyll *a*

The KN210-04 cruise was equipped with a SBE9+ CTD with a depth limit of 6000m. We used a SBE3T/SBE4C sensor system to measure temperature and conductivity and a Wet Labs FLNTURTD combination fluorometer and turbidity sensor to detect fluorescence. We calibrated fluorescence data with direct chlorophyll *a* measurements from Gwenn Hennon (Lamont Doherty Earth Observatory) determined according to methods from Arar & Collins (1997). The AE1409 cruise had a SBE 9/11 Plus CTD with a depth limit of 6800 m with a SBE 3plus temperature sensor and a SBE 4C conductivity sensor. A Chelsea Aquatracka II was used to measure fluorescence. This fluorescence was not calibrated and so is reported only in relative amounts.

### Calculations and computational tools

We primarily used MATLAB (R2014a; MathWorks, Natick, MA) to process data and create figures. The “corr” function was used to perform Spearman and Pearson correlations. The “boxplot” function was used for creation of boxplots. We performed non-metric multidimensional scaling (NMS) and accompanying statistical analyses including permutational multivariate analysis of variance (PERMANOVA), multivariate analysis of variance (MANOVA), and analysis of variance (ANOVA) to compare within group variance generated by the “betadisper” function with the community ecology package (vegan) in R (R Core Team 2015; Oksanen et al. 2017). We created the map and supplemental profile images with Ocean Data View (Schlitzer 2016).

We calculated metabolite fluxes into the particle traps with the following equation:

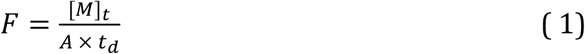

where F is the flux (nmol m^−2^ d^−1^), [M]_t_ is the abundance of a metabolite M captured in the net trap (nmol), A is the area of trap opening (1.23 m^2^), and td is the trap deployment time (days). We calculated the steady-state export turnover rates (TR) of metabolites measured both at the surface and in the net trap at a given station using:

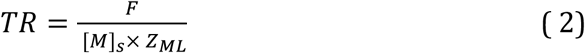

where TR is the export turnover rate (d^−1^), F is the flux from Equation 1 (nmol m^−2^ d^−1^), [M]_s_ is the concentration of metabolite M in the surface (nmol m^−3^), and Z_ML_ is the mixed layer depth (m). Durkin et al. (2016) calculated Z_ML_ for the KN210-04 stations as the depth at which downcast temperatures decreased more than 0.15 °C m^−1^; and we used the same definition for Z_ML_ at AE1409 stations (see Table S1 for mixed layer depth at each station). The export turnover rate is the fraction of the mixed layer inventory of a molecule that is being exported based on the flux of the metabolite at 150 m. The mixed layer inventory is calculated by assuming that the surface concentration of the metabolite is constant throughout the mixed layer.

### Small subunit ribosomal RNA gene sequencing

We collected samples for high resolution bacterial and archaeal small subunit ribosomal RNA (SSU or 16S rRNA) gene amplicon sequencing by filtering 2 liters of seawater through a 0.2 *μ*m Sterivex filter unit (Millipore) with an in-line GF/D (nominally 2.7 *μ*m pore size) prefilter. Following filtration, we immediately froze the Sterivex filter at −80 °C until processing. We extracted the genomic DNA from Sterivex filters as described in Hawley et al. (2017) and we used the DNA to generate Illumina 16S rRNA gene sequences (iTags) by PCR amplification of the V4-V5 hypervariable region at the DOE Joint Genome Institute (JGI). Samples were sequenced according to the standard operating protocol on an Illumina MiSeq platform at JGI (Rivers 2016). 36,361,358 iTag sequences were returned, which were binned into 4940 OTU clusters using a 97% cutoff. Further details on the 16S rRNA gene sequencing methods are provided in the supplementary information.

## Results

### Sample types

We collected samples along a latitudinal transect in the Western Atlantic Ocean from 20 °N to 38 °S. This encompassed ocean regions including the South Atlantic Subtropical Gyre, the equatorial region, and the tropical North Atlantic, including the Amazon River plume (Fig. 1). These different oceanic regions could be distinguished on the basis of their salinity profiles (Fig. S1). At eleven stations, we measured a targeted set of metabolites in sinking particles from 150 m and in suspended particles from the surface, deep chlorophyll maximum (DCM), and, in some cases, at 250 m (see Fig. S2 for chlorophyll *a* profile defining the DCM). At six additional stations, we measured metabolites in either sinking particles (2 stations) or suspended particles only (4 stations; Fig. 1). All reported metabolite concentrations are the sum of metabolites observed on all filters at a specific depth, thus representing metabolite concentrations captured on particles greater than 0.2 *μ*m. Collectively, this dataset provides context to compare overall compositional differences between suspended and sinking particles across distinct oceanic regions.

**Figure 1.**
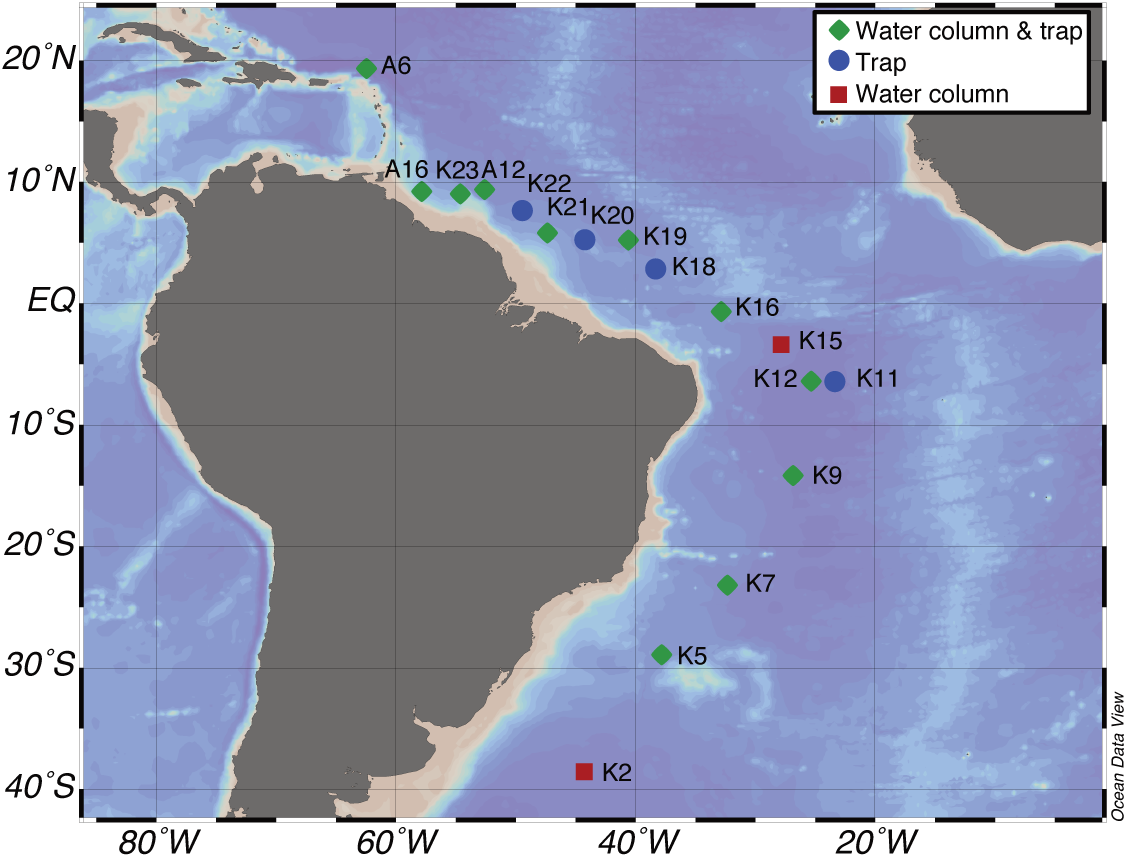
Map of transect. Station marker color and shape designates sample types.

### Metabolite composition of sinking particles

To compare the targeted metabolite compositions of sinking particles collected along the transect, we calculated the flux of each metabolite by integrating the concentration of the metabolite over the depth of the mixed layer and normalizing to the deployment duration of the particle traps. This resulted in flux values (F) of nmol m^−2^ d^−1^ for each detected metabolite. After data quality controls, we considered 34 metabolites for further study, of which 27 metabolites occurred in the sinking particle samples (Table 1; (see Johnson et al. 2017 for detection limits)). We selected the 19 most abundant metabolites (each with a flux of at least 1 nmol m^−2^ d^−1^ at one or more stations) to focus on the measured metabolites that contribute most significantly to the flux of small molecules in these sinking particles. This flux included metabolite classes such as osmolytes, amino acids, and nucleotides (Fig. 2). The total flux of these 19 metabolites into the particle traps at 150 m showed some variability along the transect. However, glycine betaine and dimethylsulfoniopropionate (DMSP) dominated all the sinking particle samples (Table 1; Fig. 2).

**Table 1.** Flux of metabolites (nmoles m^−2^ d^−1^) on sinking particles collected into the net trap deployed at 150 m. These metabolites passed quality control checks in the net trap samples. n = the number of net trap samples in which each metabolite was detected. nd = the metabolite was not detected.

| Compound | n | Station |  |  |  |  |  |  |  |  |  |  |  |  |  |  |
| --- | --- | --- | --- | --- | --- | --- | --- | --- | --- | --- | --- | --- | --- | --- | --- | --- |
|  |  | K5 | K7 | K9 | K11 | K12 | K16 | K18 | K19 | K20 | K21 | K22 | K23 | A16 | A12 | A6 |
| HBA<sup>a</sup> | 2 | 0.62 | nd | nd | nd | nd | 1.5 | nd | nd | nd | nd | nd | nd | nd | nd | nd |
| MTA<sup>b</sup> | 13 | 0.027 | 0.021 | 0.033 | 0.036 | 0.049 | 0.048 | 0.044 | 0.048 | 0.061 | 0.037 | 0.063 | 0.018 | 0.052 | nd | nd |
| Adenosine | 15 | 3.9 | 4.0 | 3.9 | 12 | 4.0 | 8.2 | 6.5 | 6.8 | 7.9 | 11 | 3.7 | 1.7 | 11 | 14 | 49 |
| AMP<sup>c</sup> | 5 | nd | nd | nd | nd | nd | nd | 0.49 | 6.2 | 0.90 | nd | nd | 0.23 | nd | nd | 1.1 |
| Arginine | 4 | nd | 1.3 | nd | nd | nd | 3.0 | 2.5 | nd | nd | nd | nd | 1.2 | nd | nd | nd |
| Biotin | 2 | nd | nd | nd | nd | 0.19 | nd | nd | nd | nd | nd | nd | 0.012 | nd | nd | nd |
| Caffeine | 12 | 3.0 | 2.6 | 4.8 | 5.0 | 5.1 | 5.0 | 5.1 | 2.8 | 4.2 | 3.1 | 4.7 | 2.4 | nd | nd | nd |
| Choline | 15 | 23 | 19 | 40 | 33 | 51 | 32 | 52 | 31 | 58 | 40 | 34 | 11 | 30 | 22 | 51 |
| Citric acid | 5 | nd | nd | 6.7 | nd | nd | nd | nd | nd | nd | nd | nd | 15 | 3.7 | 2.8 | 18 |
| Cytosine | 3 | nd | nd | nd | 4.7 | nd | nd | nd | nd | nd | nd | nd | nd | 4.7 | nd | 33 |
| DMSP<sup>d</sup> | 15 | 86 | 120 | 19 | 180 | 60 | 180 | 55 | 62 | 48 | 52 | 53 | 42 | 82 | 220 | 220 |
| Folic acid | 1 | nd | nd | nd | nd | nd | nd | nd | 0.070 | nd | nd | nd | nd | nd | nd | nd |
| Glycine betaine | 15 | 260 | 230 | 450 | 420 | 380 | 480 | 410 | 270 | 310 | 260 | 380 | 110 | 210 | 280 | 560 |
| Guanine | 15 | 72 | 93 | 270 | 150 | 100 | 42 | 61 | 33 | 72 | 31 | 32 | 21 | 69 | 67 | 230 |
| Indole 3-acetic acid | 10 | nd | 0.020 | 0.034 | 0.040 | 0.058 | 0.022 | 0.028 | 0.031 | 0.049 | 0.016 | nd | nd | nd | nd | 0.27 |
| Inosine | 14 | 3.7 | 1.5 | 1.6 | 2.3 | 5.5 | 1.6 | 4.0 | 8.0 | 4.9 | 2.6 | 1.5 | 0.32 | nd | 2.0 | 4.6 |
| (Iso)leucine | 14 | 3.2 | 4.9 | 3.4 | 1.8 | 4.0 | 4.2 | 12 | 14 | 11 | 8.3 | 4.5 | 1.6 | 1.9 | nd | 21 |
| Malic acid | 2 | nd | nd | 5.6 | nd | nd | nd | 4.8 | nd | nd | nd | nd | nd | nd | nd | nd |
| Methionine | 8 | 5.1 | 0.33 | nd | 1.2 | nd | nd | nd | 13 | 9.4 | 9.6 | nd | nd | 3.8 | nd | 34 |
| Pantothenic acid | 1 | nd | nd | nd | nd | nd | nd | nd | nd | 0.38 | nd | nd | nd | nd | nd | nd |
| Phenylalanine | 15 | 2.2 | 1.8 | 1.4 | 2.7 | 2.0 | 4.1 | 3.8 | 13 | 9.7 | 6.4 | 4.3 | 1.0 | 3.2 | 2.1 | 15 |
| Proline | 13 | 7.5 | 52 | 27 | 28 | 37 | 130 | 53 | 20 | 23 | 45 | 22 | nd | nd | 20 | 73 |
| Pyridoxine | 1 | nd | nd | nd | 0.25 | nd | nd | nd | nd | nd | nd | nd | nd | nd | nd | nd |
| Riboflavin | 14 | 0.031 | 0.043 | 0.021 | 0.040 | 0.055 | 0.11 | 0.071 | 0.061 | 0.13 | 0.15 | 0.057 | 0.021 | 0.089 | nd | 0.11 |
| Taurocholate | 4 | 0.050 | nd | nd | nd | nd | nd | nd | 0.39 | 0.10 | 0.48 | nd | nd | nd | nd | nd |
| Tryptophan | 15 | 1.0 | 1.2 | 1.2 | 1.5 | 1.7 | 2.1 | 2.7 | 9.5 | 7.7 | 3.8 | 2.7 | 0.65 | 1.3 | 1.8 | 4.8 |
| Xanthine | 15 | 3.4 | 7.4 | 10 | 7.9 | 14 | 5.7 | 19 | 14 | 43 | 7.0 | 7.7 | 8.8 | 2.2 | 2.1 | 6.9 |
| 2,3-Dihydroxybenzoic acid | 0 | nd | nd | nd | nd | nd | nd | nd | nd | nd | nd | nd | nd | nd | nd | nd |
| 3-Mercaptopropionate | 0 | nd | nd | nd | nd | nd | nd | nd | nd | nd | nd | nd | nd | nd | nd | nd |
| 4-Aminobenzoic acid | 0 | nd | nd | nd | nd | nd | nd | nd | nd | nd | nd | nd | nd | nd | nd | nd |
| Desthiobiotin | 0 | nd | nd | nd | nd | nd | nd | nd | nd | nd | nd | nd | nd | nd | nd | nd |
| IMP<sup>c</sup> | 0 | nd | nd | nd | nd | nd | nd | nd | nd | nd | nd | nd | nd | nd | nd | nd |
| NAD<sup>f</sup> | 0 | nd | nd | nd | nd | nd | nd | nd | nd | nd | nd | nd | nd | nd | nd | nd |
| TMP<sup>g</sup> | 0 | nd | nd | nd | nd | nd | nd | nd | nd | nd | nd | nd | nd | nd | nd | nd |

<sup>a</sup>4-hydroxybenzoic acid, <sup>b</sup>5-methylthioadenosine, <sup>c</sup>adenosine 5'-monophosphate, <sup>d</sup>dimethylsulfonylpropionate, <sup>e</sup>inosine 5'-monophosphate, <sup>f</sup>nicotinamide adenine dinucleotide, <sup>g</sup>thiamin monophosphate

**Figure 2.**
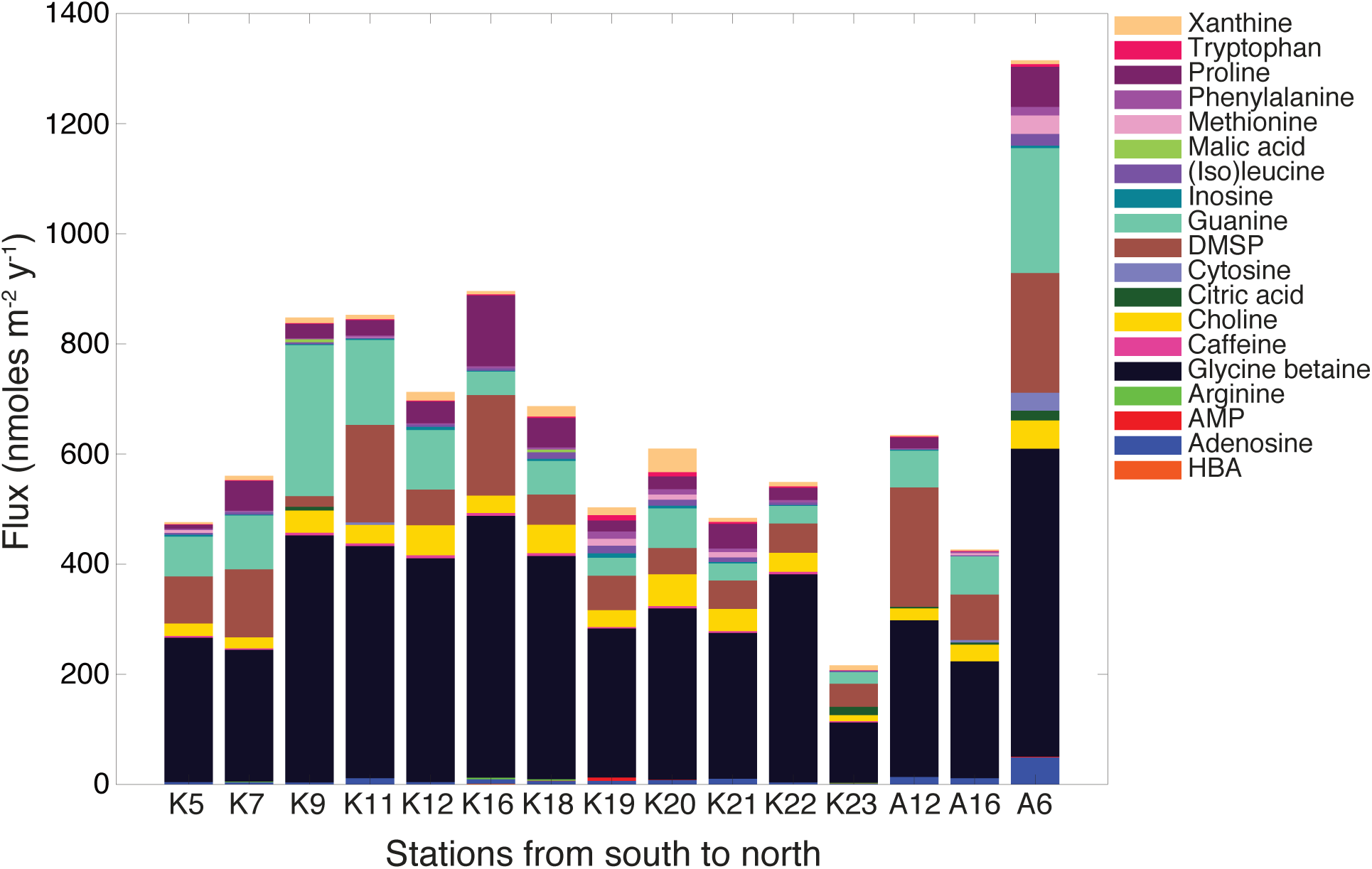
Fluxes of metabolites on the sinking particles collected in the net traps at 150 m. Only metabolites with a flux greater than 1 nmol m^−2^ d^−1^ in at least one sample are shown. K indicates samples from cruise KN210-04 whereas A indicates samples from cruise AE1409. Stations are presented in order of latitude from south (left) to north (right).

At all stations, the flux of the osmolyte glycine betaine was the largest of any metabolite, ranging from 110–560 nmoles m^−2^ d^−1^. Other sinking particle metabolite fluxes were more variable. For example, guanine was a larger component of flux (270 nmoles m^−2^ d^−1^) into the net trap at station K9 while DMSP and proline contributed more to the flux (180 and 130 nmoles m^−2^ d^−1^, respectively) at station K16 (Fig. 2). Choline was a sizable component of the metabolite flux at many stations ranging from 11–58 nmoles m^−2^ d^−1^, in some cases comparable to the contribution of DMSP which ranged from 19–220 nmoles m^−2^ d^−1^. Other metabolites such as adenosine, caffeine, inosine, (iso)leucine, phenylalanine, riboflavin, tryptophan, and xanthine contributed much smaller total moles to the flux of these metabolites but were measured throughout the transect (Table 1). In total, the described metabolites collectively contributed a small fraction of carbon in sinking particle organic matter (<0.4% by weight; Fig. S3).

### Comparison of metabolite composition of suspended and sinking particles

Our primary motivation was to compare the metabolite composition of suspended particles to that of sinking particles to determine how metabolite composition changes as particles sink. We measured most metabolites in both the suspended particles and the sinking particles (Table 2). Only three metabolites (pantothenic acid, cytosine, and indole 3-acetic acid (IAA)) were detected exclusively in the sinking particles. These metabolites (a vitamin, a nucleobase and a signaling molecule, respectively) are usually in lower abundance within cellular biomass relative to other metabolites (Bennett et al. 2009) as might be expected due to their cellular roles. Thus, it is likely that we detected these metabolites only in the sinking particles because there was more biomass in general in these samples than in the suspended particle samples. The summed total of targeted metabolites collected was, on average, approximately 6x more in the sinking particle samples than in the suspended particle samples (Fig. S4).

**Table 2.** Number of samples at each water column depth and in the particle traps that contained each metabolite. In parentheses the percentage of the total samples in each category that contained the metabolite. n = the number of samples in each category

| **Compound** | **Surface**<br>**n = 14** | **DCM**<br>**n = 17** | **250 m**<br>**n = 7** | **Particle traps**<br>**n = 15** |
| --- | --- | --- | --- | --- |
| 2,3-Dihydroxybenzoic acid | 0 (0%) | 0 (0%) | 0 (0%) | 0 (0%) |
| 3-Mercaptopropionate | 0 (0%) | 0 (0%) | 0 (0%) | 0 (0%) |
| 4-Aminobenzoic acid | 0 (0%) | 0 (0%) | 0 (0%) | 0 (0%) |
| HBA<sup>a</sup> | 2 (14%) | 0 (0%) | 0 (0%) | 2 (13%) |
| MTA<sup>b</sup> | 11 (79%) | 13 (76%) | 0 (0%) | 13 (87%) |
| Adenosine | 11 (79%) | 13 (76%) | 3 (43%) | 15 (100%) |
| AMP<sup>c</sup> | 11 (79%) | 16 (94%) | 6 (86%) | 5 (33%) |
| Arginine | 2 (14%) | 2 (12%) | 4 (57%) | 4 (27%) |
| Biotin | 2 (14%) | 1 (6%) | 1 (14%) | 2 (13%) |
| Caffeine | 4 (29%) | 8 (47%) | 6 (86%) | 12 (80%) |
| Cytosine | 0 (0%) | 0 (0%) | 0 (0%) | 3 (20%) |
| Desthiobiotin | 0 (0%) | 0 (0%) | 0 (0%) | 0 (0%) |
| DMSP<sup>d</sup> | 14 (100%) | 17 (100%) | 2 (29%) | 15 (100%) |
| Folic acid | 1 (7%) | 0 (0%) | 0 (0%) | 1 (7%) |
| Glycine betaine | 13 (93%) | 17 (100%) | 2 (29%) | 15 (100%) |
| Guanine | 14 (100%) | 17 (100%) | 7 (100%) | 15 (100%) |
| Indole 3-acetic acid | 0 (0%) | 0 (0%) | 0 (0%) | 10 (67%) |
| Inosine | 11 (79%) | 8 (47%) | 3 (43%) | 14 (93%) |
| IMP<sup>c</sup> | 0 (0%) | 1 (6%) | 0 (0%) | 0 (0%) |
| (Iso)leucine | 7 (50%) | 10 (59%) | 3 (43%) | 14 (93%) |
| Methionine | 0 (0%) | 1 (6%) | 2 (29%) | 8 (53%) |
| Pantothenic acid | 0 (0%) | 0 (0%) | 0 (0%) | 1 (7%) |
| Phenylalanine | 14 (100%) | 17 (100%) | 7 (100%) | 15 (100%) |
| Proline | 11 (79%) | 8 (47%) | 5 (71%) | 13 (87%) |
| Pyridoxine | 3 (21%) | 3 (18%) | 3 (43%) | 1 (7%) |
| Riboflavin | 2 (14%) | 4 (24%) | 0 (0%) | 14 (93%) |
| Tryptophan | 13 (93%) | 17 (100%) | 6 (86%) | 15 (100%) |
| Xanthine | 5 (36%) | 9 (53%) | 0 (0%) | 15 (100%) |

<sup>a</sup>4-hydroxybenzoic acid, <sup>b</sup>5-methylthioadenosine, <sup>c</sup>adenosine 5'-monophosphate, <sup>d</sup>dimethylsulfoniopropionate, <sup>e</sup>inosine 5'-monophosphate

To facilitate comparisons between the overall metabolite composition of the suspended and sinking particles we calculated the mole fraction of each metabolite per sample (M_i_/M_T_, where M_i_ is the concentration of metabolite i and M_T_ is the total concentration of metabolites in that sample; Table S3). To distinguish data normalized in this way from data reported as a concentration or flux we refer to these data as the relative targeted metabolite composition of a sample.

When the relative targeted metabolite compositions of the samples were compared using non-metric multidimensional scaling, three overlapping clusters emerged (Fig. 3). Suspended particles from the surface and DCM formed a scattered cluster that was not clearly distinct from the suspended particles collected at 250 m and the sinking particles. However, the sinking particles formed a tight cluster, highlighting the similarity of their metabolite composition. Analysis of variance (ANOVA) of the variance of the sinking particle composition compared to the suspended particle composition (using the betadisper function in the vegan package in R) indicates that the sinking particles were significantly more homogenous than the suspended particles (*F* = 13.7, *p* = 0.0005, df = 1). A PERMANOVA indicated significant differences between the suspended and sinking particles (*p* < 0.001, df = 1) but the high variability in the suspended particle composition resulted in a low *R^2^* value of 0.30. The observed pattern of clustering was driven by the relative amounts of metabolites in these samples. The length of each arrow (Fig. 3) indicates the strength of the relationship between the relative abundance of a metabolite and the sample cluster (*p* < 0.0001 for all shown metabolites). For instance, DMSP was a larger mole fraction of suspended particles in the surface and DCM, while phenylalanine and guanine comprised more of the metabolite pool in deeper suspended particle samples around 250 m. Glycine betaine, xanthine, and proline were larger mole fractions in the sinking particles than in the suspended particles.

**Figure 3.**
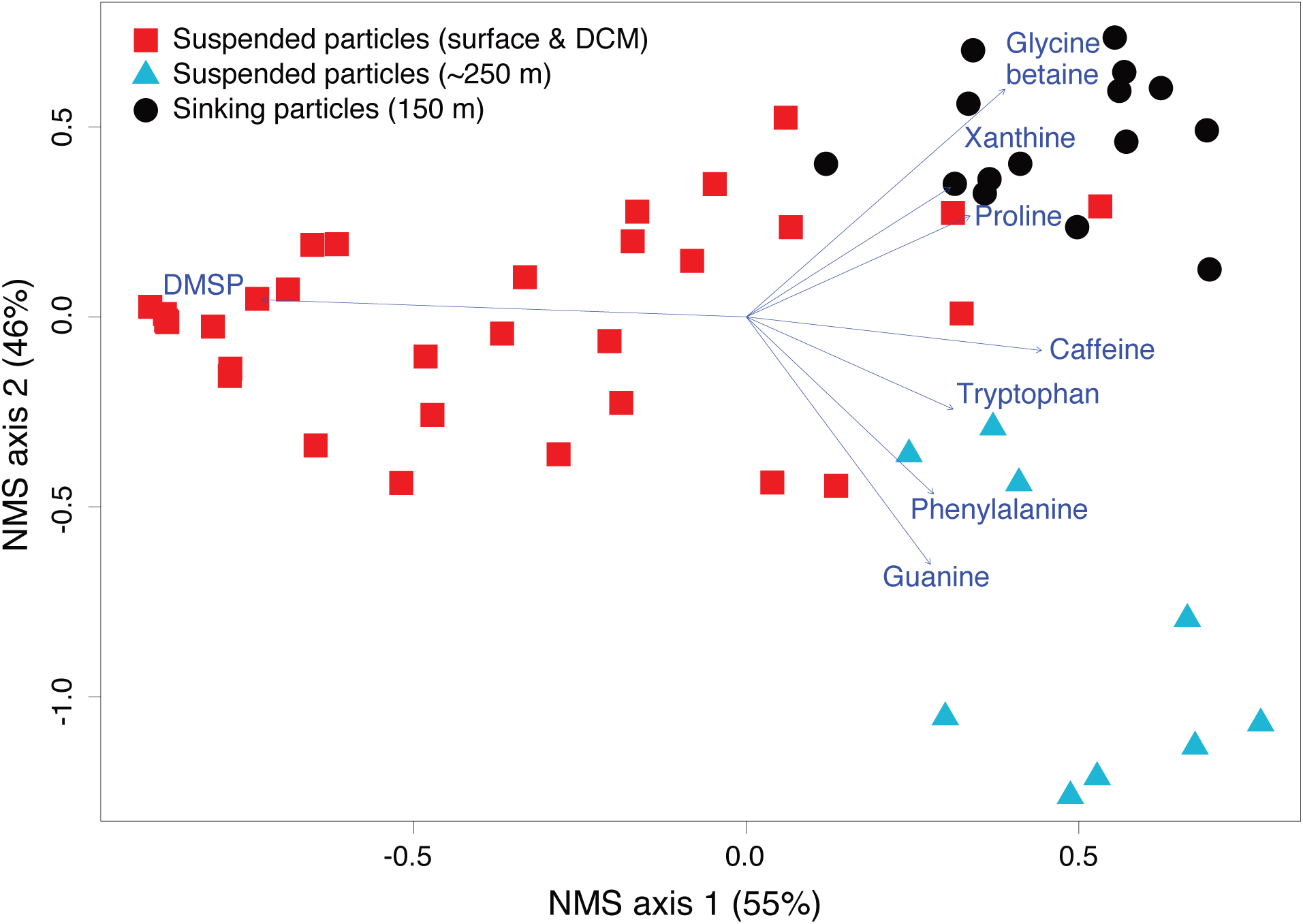
Non-metric multidimensional scaling (NMS) shows that the metabolite compositions of the suspended particles are distinct from that of the sinking particles. The stress on the NMS is 0.0584. Each marker represents a different sample and they are clustered by the relative composition of the metabolites in each sample. The length of the lines for each metabolite indicate the strength of a relationship between a metabolite and a sample cluster (i.e. a longer line indicates a stronger relationship).

Similarly, a significant difference in microbial community composition between sinking particles and suspended particles was evident in complementary microbial diversity data from these samples (Fig. S5, NPMANOVA, *p* = 0.0010, df = 1) and consistent with prior results (DeLong et al. 1993; Fontanez et al. 2015).

To examine whether there was a relationship between the metabolite distributions observed in the sinking particles and their source material in the euphotic zone, several attributes of the source particles were compared to the metabolite composition of the sinking particles. The total flux of targeted metabolites did not have a significant relationship with the total particulate organic carbon (POC) flux (Fig. 4a), and the abundance of an individual metabolite on the sinking particles showed no relationship to its abundance on the euphotic zone suspended particles (i.e. the source particles) (Fig. 4b; see Table S4 for Spearman and Pearson correlation results that support the null hypothesis). Although metabolites that were higher in abundance in the surface suspended particles were also higher in abundance in the sinking particles, this is to be expected due to their physiological roles in cells (i.e. osmolytes like DMSP and glycine betaine are high and pathway intermediates and vitamins like methylthioadenosine (MTA) and riboflavin are low). The type of particles that were numerically influential in the overall sinking particle composition based on principal components analysis (PCA) (diatoms, coccolithophores, dinoflagellates, nanocells, or aggregates as determined in Durkin et al. 2016) at the different stations also did not overall appear to influence the metabolite composition (Fig. 4c). NMS of the sinking particle samples based on their metabolite composition showed no clustering and no relationship between the sinking particle metabolite composition and the main particle types that composed the sinking particle pool. However, the average flux of dinoflagellates into the traps was significantly positively correlated with the flux of DMSP along the transect (Spearman correlation: *rho* = 0.88, *p* = 0.008, df = 5; Table S5). Otherwise, we found no significant relationship between the metabolite composition of sinking particles and the source material.

**Figure 4.**
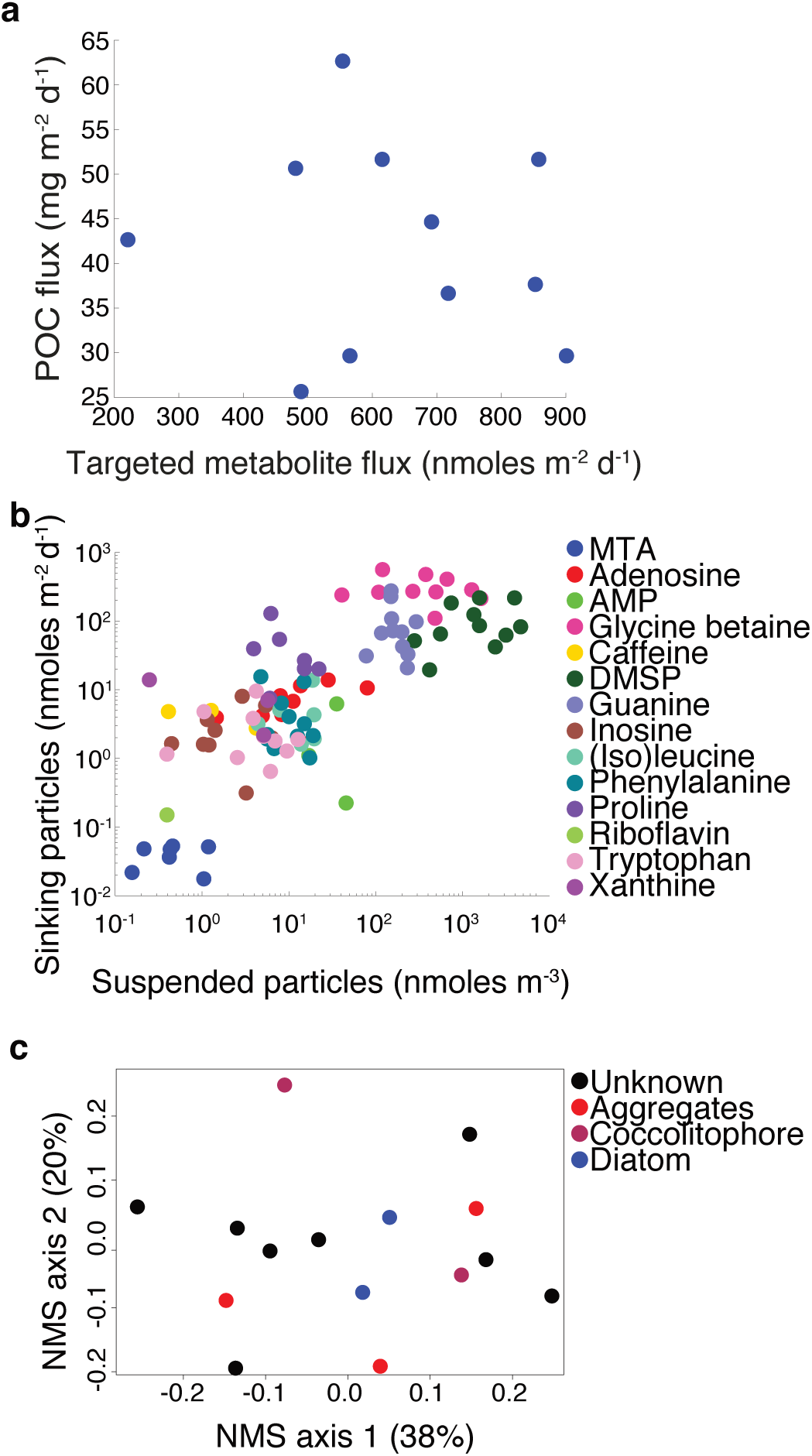
Comparison of metabolite flux and composition to possible factors that may influence metabolite composition and abundance on sinking particles. **a)** Flux of targeted metabolites compared to the particulate organic carbon (POC) flux. **b)** Comparison of metabolite abundance in surface suspended particles versus the flux of metabolites on the sinking particles. **c)** Nonmetric multidimensional scaling (NMS) plot of relative metabolite composition of trap samples. Markers are colored based on dominant particle type assigned to that station by Durkin et al. (2016). NMS Stress = 0.173.

### Metabolite production and removal on sinking particles

As particles sink, metabolites derived from the surface ocean will be removed while other metabolites will appear through production by particle-associated microbial communities through either *de novo* synthesis or degradation of organic matter. To infer the extent to which individual metabolite abundances were controlled by production or removal on the sinking particles, the export turnover rate, or rate of export out of the mixed layer, was calculated. For this study, we built on the work of Durkin et al. (2016) who calculated export turnover rates for phytoplankton cells for stations K7, K9, K12, K16, K19, K21, and K23. Included in this calculation were the surface abundances of diatoms, coccolithophores, dinoflagellates, and nanocells (referring to all small cells (~5 *μ*m) that can be observed with 400★ magnification under a microscope).

We calculated the export turnover rate for each metabolite where we had simultaneous measurements within sinking and suspended particles. Overall, this calculation could be completed at the eleven stations where water column and particle trap data were both available (stations K5, K7, K9, K12, K16, K19, K21, K23, A16, A12, and A6; shown in green, Fig. 1).The export turnover rates of metabolites ranged from 6 × 10^−5^ d^−1^ – 6 × 10^−1^ d^−1^ (geometric mean: 7 × 10^−3^ d^−1^) (Table S6; Fig. 5). In comparison, phytoplankton cells had export turnover rates that were generally lower ranging from 2 × 10^−6^ d^−1^ – 3 × 10^−3^ d^−1^ (geometric mean: 8 × 10^−5^ d^−1^) (Durkin et al. 2016).

**Figure 5.**
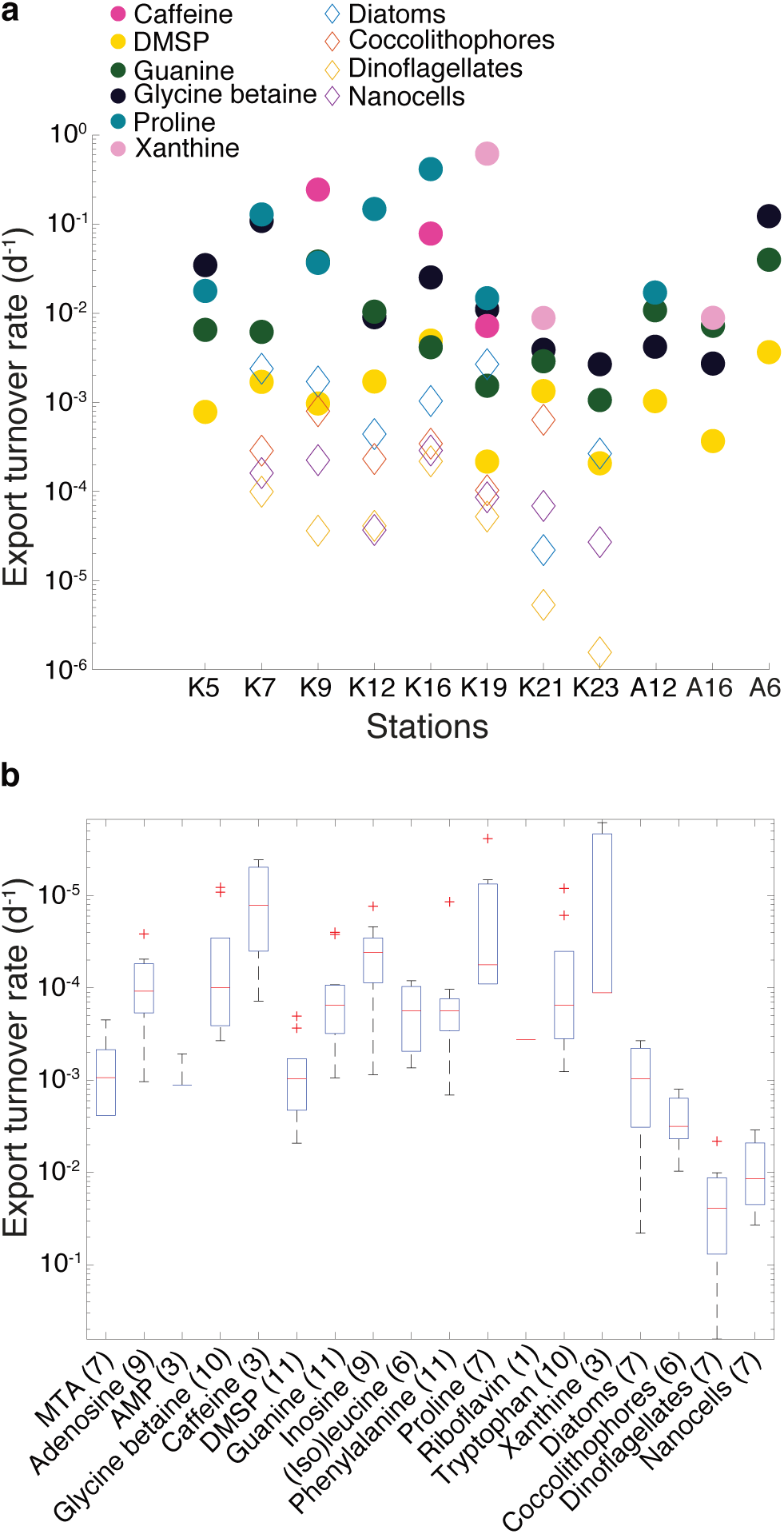
a) The export turnover rates for selected metabolites that were detected in both the suspended particles in the surface and in the sinking particles for a given station. The export turnover rates of phytoplankton cells are plotted with open diamond markers (obtained from Durkin et al. 2016). **b)** The range of export turnover rates for each metabolite and phytoplankton cells from all stations. The red line indicates the median value, the bottom blue line is the 25^th^ percentile, the top blue line is the 75^th^ percentile, the whiskers encompass all non-outliers, and red (+) indicates outliers. In parentheses, the number of export turnover rates that were calculated for each metabolite.

Because metabolites on sinking particles might also be derived from the DCM (all particle traps (150 m) were below the DCM), the same export turnover rate calculation was performed using the metabolite abundances from the DCM (Fig. S6). Generally, the trends remained the same with the exception of guanine, which has a lower export turnover rate when calculated at the DCM. This is expected because guanine abundances are greater at the DCM than at the surface.

The phytoplankton export turnover rates provided a baseline comparison as their source is constrained to the euphotic zone, while the metabolite export turnover rates reflect a spectrum of production versus removal processes on the timescale of the sinking particles (Fig. 5). For metabolites that are only removed during transport (and not produced), we would predict export rates that were similar to those of intact phytoplankton cells. In contrast, metabolites actively being produced on the sinking particles would have greater rates of export from the mixed layer than intact phytoplankton cells or metabolites that were exclusively being removed. We considered whether higher export rates of certain metabolites could be derived from associations with heavier particles like fecal pellets or aggregates that sink more rapidly than phytoplankton cells, thus removing metabolites from the euphotic zone more quickly. However, faster sinking detrital material, which was a major component of the sinking particle samples (Durkin et al. 2016), will be more heavily colonized with heterotrophic microbes (Ploug and Grossart 2000) than individual phytoplankton cells, thus these aggregates are where most active production of metabolites would be occurring. Thus, our hypothesis that metabolites with higher export rates are indicative of active production of that molecule on the sinking particles should be relatively independent of sinking speed in this dataset.

DMSP had the lowest turnover rates of any metabolite in eight of the eleven stations for which this calculation was performed. This suggests that a lower fraction of surface-produced DMSP reaches 150 m through transport on these sinking particles compared to other metabolites. In contrast, the metabolites with the highest export turnover rates vary by station but include proline, inosine, caffeine, adenosine, and glycine betaine (Fig. 5; Table S6).

Similarly, the differences in metabolite abundances from euphotic zone suspended particles versus sinking particles can be seen by plotting the mole fraction of a metabolite along the transect (Fig. 6). For instance, DMSP reached upwards of 80% of the total moles of targeted metabolites measured in the surface and DCM suspended particles (Fig. 6a). In contrast, though present in the sinking particles, DMSP was only ~20-30% of the total moles of targeted metabolites measured. Inversely, glycine betaine reached between 40–60% of the total composition of the sinking particles while being a smaller component in the suspended particles (Fig. 6b). Proline, an amino acid, and xanthine, a nucleotide derivative, also represented higher mole fractions on the sinking particles than on the suspended particles (Fig. 6c-d).

**Figure 6.**
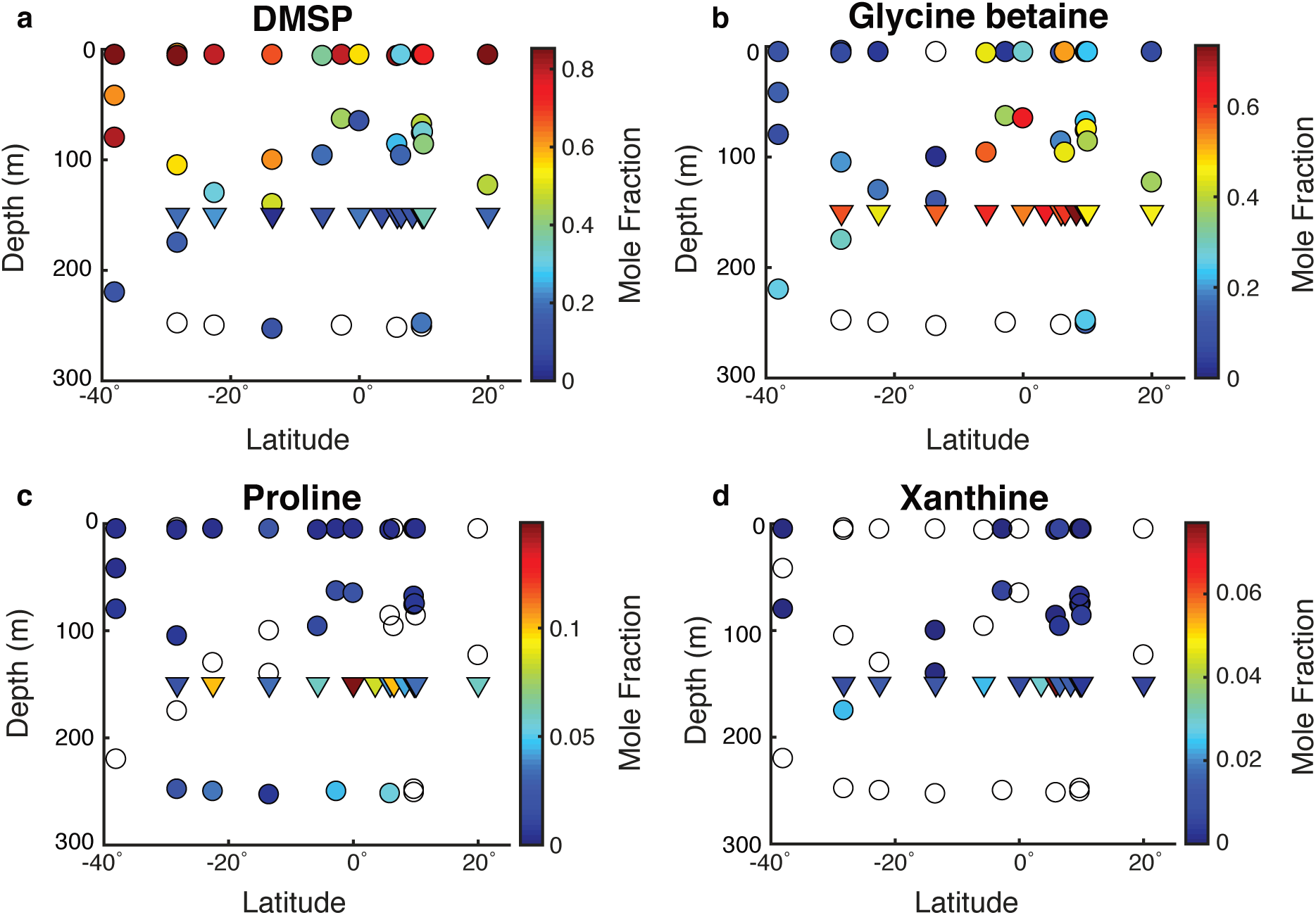
Metabolite profiles in both the suspended (circles) and sinking particles (triangles). Open symbols indicate samples where the metabolite was not detected. Warmer colors indicate a higher mole fraction of that metabolite (metabolites normalized to total moles targeted metabolites) in the sample. Shown are **a)** DMSP, **b)** glycine betaine, **c)** proline, **d)** xanthine.

## Discussion

### Metabolite composition of sinking particles driven by colonizing microbial community rather than by euphotic zone source material

The relative metabolite composition of sinking particles was distinct from that of suspended particles. The metabolite composition of the sinking particles showed no relationship to the overall POC flux, the metabolite composition of the source particles (i.e. the suspended euphotic zone particles), or to whether diatom cells, coccolithophore cells, or aggregates were numerically important components of the sinking particles (Fig. 4). When comparing the fluxes of individual metabolites directly to the flux of particle types, there was a positive correlation along the transect between the flux of DMSP and the flux of dinoflagellate cells (Table S6).

While dinoflagellates cells would not be the only source of DMSP, they would certainly be an expected source of DMSP to the sinking particle community. Thus, while overall metabolite composition of the sinking particles does not seem to be driven by factors that shaped the sinking particles in the surface ocean, the abundance of an exclusively surface ocean-derived metabolite like DMSP does reflect its potential source. In contrast, the distributions of other metabolites that are likely derived from the metabolic activity of the sinking particle-associated microbial community suggests a functionally similar microbial community throughout the transect.

The dominant metabolic pathways required for degradation of the organic matter on sinking particles differ from those required by free-living, euphotic zone-dwelling microbes (Fontanez et al. 2015), and thus are likely to be reflected in distinct metabolic profiles. The microbial diversity data supports this hypothesis, showing that the prokaryotic community composition on the sinking particles differed from that of the suspended particles (Fig. S5). Moreover, the differences between communities on sinking particles across the transect were generally smaller than the differences between communities on suspended particles at a single station. Similar bacterial communities on sinking particles could lead to similar metabolite compositions, particularly if the metabolic pathways associated with organic matter degradation produce a characteristic profile of metabolic intermediates. This composition could be dynamic reflecting a succession process as the particles are transformed on their downward transport through the water column (Pelve et al. 2017). We expect that the metabolite composition of the sinking particles along the transect is shaped by the removal of metabolites through microbial respiration, the production of molecules from the breakdown of larger molecules derived from the sinking particle organic matter pool, and production of molecules associated with the physiological requirements of the sinking particle-associated microbial community. In the following sections examples of metabolite distributions potentially shaped by these factors will be illustrated. However, more work with sinking particles collected at multiple depths is required to assess these hypotheses and to better constrain the particle niche space in relation to biogeochemical fluxes in the ocean.

### Osmolytes as indicators of microbial population dynamics and physiological changes associated with particle colonization

Osmolytes are predominantly organic molecules present in the cytosol of cells that assist in the regulation of osmotic pressure as well as contribute to maintaining protein structure (Yancey et al. 1982). Due to their high concentrations within cells (~millimolar cytosolic concentrations), they tend to be especially abundant metabolites. However, their abundances can shift in response to external changes in temperature, salinity, and pressure, which dictate the osmotic stress the cell experiences (Yancey et al. 2002; Yancey 2005; Gebser and Pohnert 2013), and in response to the availability of osmolytes in the environment (Burg and Ferraris 2008). Additionally, some osmolytes may be exclusively used by certain groups of microbes. For example, the osmolyte DMSP is exclusively an algal metabolite in the open ocean (Stefels 2000). Thus, the fact that we observed higher mole fractions of DMSP in euphotic zone suspended particle samples than in sinking particles (Fig. 3; Fig. 6a) is consistent with its origins as an algal metabolite. By contrast, in sinking particles, glycine betaine, a ubiquitous osmolyte, was a larger mole fraction than DMSP (Fig. 3; Fig. 6b).

We hypothesize that this compositional shift from DMSP as the dominant osmolyte to glycine betaine is due to both the degradation of DMSP as particles sink, as well as to the increased prevalence of heterotrophic microbes that synthesize glycine betaine but not DMSP. Interestingly, in *Escherichia coli* glycine betaine acts as an osmoprotectant under both aerobic and anaerobic conditions, while its precursor, choline, only acts as an osmoprotectant under aerobic conditions (Landfald and Str0m 1986), suggesting that glycine betaine may be more extensively used as an osmolyte on sinking particles where anoxic zones exist (Bianchi et al. 2018). The export turnover rates lend further support and nuance to this idea, indicating that DMSP has similar export turnover rates to those of the phytoplankton cells that would produce it, while glycine betaine has higher export turnover rates than the phytoplankton cells (at stations where it is measured in the surface; Fig. 5a-b). Many heterotrophic bacteria in the ocean catabolize DMSP to obtain energy (Reisch et al. 2011) and to obtain carbon and sulfur for biomass (Simó et al. 2009); thus DMSP is likely preferentially degraded on sinking particles. We do not expect DMSP to be regenerated on these particles because the organisms that produce it would be dead or metabolically inactive when they exit the euphotic zone. In contrast, glycine betaine is utilized as an osmolyte by organisms across the domains of life, although it is less commonly used by phytoplankton than DMSP (Yancey et al. 1982; Boch et al. 1997; Keller et al. 1999; Lai and Lai 2011). Thus, on sinking particles, glycine betaine could be catabolized (González et al. 1999) but is likely also produced by resident particle microbial communities.

The amino acid proline was a larger mole fraction of the sinking particles than the suspended particles and contributed more to the sinking particle flux than any other measured amino acid (Fig. 2). At station K16, proline was a larger component of the flux than guanine or choline, contributing around 10% of the total detected moles. The proline export turnover rate at a number of stations was among the higher rates measured for the metabolites in this study (Fig. 5a-b), implying increased production by microbes on sinking particles relative to euphotic zone suspended particles. Relatively few measurements of proline are available in the ocean, but Takasu and Nagata (2015) recently documented that proline is more abundant than other amino acids on bacteria-sized particles at 1000 m. This abundance may be due in part to its function as an osmolyte (Burg and Ferraris 2008; Brill et al. 2011; Götz et al. 2018) and cryoprotectant (Yancey 2005). The apparent use of proline by some organisms in response to increased osmotic pressure and its greater prevalence at depth in bacteria-sized particles suggests that proline may be an important osmolyte for microbes adapted to deeper ocean regions.

### Metabolite dynamics on sinking particles as indicators of organic matter degradation processes

While metabolite distributions are shaped by the microbial community and its physiological requirements, the interaction of the unique organic matter-rich environment of sinking particles and microbial metabolism may also influence metabolite abundances on sinking particles. The availability of certain substrates likely influences the expression of individual metabolic pathways, and the presence or absence of intracellular metabolic intermediates and byproducts.

Glycine betaine offers a possible case-in-point. Glycine betaine was the largest mole fraction of the detected metabolites on the sinking particles, suggesting that it has a role as an osmolyte in these environments. In contrast, glycine betaine was not detected on suspended particles at 250 m (Fig. 6b), despite the ubiquity of biosynthetic capabilities for glycine betaine across taxa. This discrepancy leads us to hypothesize that the dominance of glycine betaine may be influenced by available exogenous substrates unique to the sinking particles. For example, choline is derived from the breakdown of phosphatidylcholine, a lipid known to be present in the surface ocean (Van Mooy and Fredricks 2010; Fulton et al. 2017) and is required by some bacteria as a precursor for the synthesis of glycine betaine (Boch et al. 1994). Utilizing choline obtained from phosphatidylcholine for catabolism has been documented in soil (Kortstee 1970; Tollefson and McKercher 1983), and utilizing choline obtained from their host is a known source of glycine betaine in pathogenic bacteria (Fitzsimmons et al. 2012). This scenario would suggest that lipid degradation on these particles might influence microbial adaptive strategies. In future work, tracing the source of glycine betaine present on sinking particles may yield insight not only into microbial physiology but into carbon and nitrogen cycling on these particles.

Xanthine was also more abundant on sinking particles compared to suspended particles in the euphotic zone (Fig. 3; Fig. 5) and may be linked to degradation processes on sinking particles. Xanthine is a purine and is structurally related to the nucleobase guanine. It was measured in all of the sinking particle samples but not all of the suspended particles from the surface and DCM (Fig. 6d). At station K19, the xanthine export turnover rates were higher than those of the other metabolites or phytoplankton cells, suggesting that a large fraction of xanthine from the surface reaches 150 m or that it is produced as these particles sink. Additionally, the xanthine export turnover rate calculated with the DCM metabolite abundance was the highest of any metabolite in the stations where it was calculated (Fig. S6).

Xanthine is part of a metabolic pathway that recycles components of nucleic acids and it is a possible degradation product of RNA or DNA (Rinas et al. 1995). Xanthine accumulates extracellularly in *Escherichia coli* cultures when they reach stationary phase, specifically the point when nutrients limit growth and dying cells equal dividing cells. This is attributed, in particular, to the degradation of ribosomal RNA (Rinas et al. 1995). It is reasonable to speculate, then, that elevated xanthine concentrations on sinking particles were due to the degradation of cellular material, perhaps derived from the nucleobase guanine, which was prevalent in the euphotic zone suspended particle samples. As a result, particle-associated marine microorganisms may have adapted to utilize xanthine. For example, the marine a-proteobacterium *Ruegeriapomeroyi* is adapted to a particle-associated lifestyle and can catabolize xanthine using the enzyme xanthine dehydrogenase (Moran et al. 2004; Cunliffe 2016).

## Conclusions

Here we show that the metabolite composition of sinking particles throughout the western South Atlantic gyre, equatorial region, and tropical North Atlantic was relatively constant and showed no relationship to the metabolite composition of suspended particles in the euphotic zone or to differences in the numerical contribution of diatom or coccolithophore cells to the overall particle flux. The observation that the metabolite composition of sinking particles differed from that of suspended particles was consistent with the observation that microbial community composition also differs on sinking particles from the community in seawater (Fontanez et al. 2015; Thiele et al. 2015; Pelve et al. 2017). This may be related to differences in community composition and metabolic strategies associated with sinking particles such as the possible utilization of proline and glycine betaine as osmolytes on sinking particles. Finally, the dynamics of individual metabolites on sinking particles relative to suspended particles suggests that certain metabolites such as xanthine and glycine betaine may be linked to organic matter degradation processes.

These findings provide a new lens through which to observe microbial community activity, the processing of organic matter on sinking particles, and ultimately carbon flux to the deep ocean. While these small molecules do not comprise the bulk of organic matter transported on sinking particles, they are the conduit through which larger organic molecules are processed. Gaining an understanding of the small molecules that dominate sinking particles will allow us to design future studies to quantitatively trace the metabolic pathways that transform and remineralize sinking organic matter. Additionally, these molecules may provide a small but important source of highly labile organic matter to deep sea environments. A recent study, for example, found that deep sea microorganisms produce transporters not only for substrates such as amino acids and carbohydrates, but also for osmolytes suggesting that they may be an important organic carbon source in the deep ocean (Bergauer et al. 2017). Understanding which molecules may be important for metabolic activity in the deep ocean is essential for understanding these difficult-to-access and low-abundance microbial populations. Future studies of these small molecules should include measuring metabolite fluxes on sinking particles along an extensive depth gradient and controlled studies of their production and utilization by marine microbes. This will allow us to determine how metabolite fluxes on sinking particles change with increasing depth, perhaps responding to differences in pressure, temperature, and substrate availability. Additional characterization of the biochemistry underlying these molecules may facilitate their use as indicators of specific globally relevant biogeochemical cycles.

## Acknowledgments

The authors would like to thank the captain and crew of the R/V *Knorr* and R/V *Atlantic Explorer*, as well as Justin Ossolinski, Catherine Carmichael, and Sean Sylva for helping to make this dataset possible. Special thanks to Colleen Durkin for sharing her data and providing feedback on the manuscript. Funding for this work came from the National Science Foundation (NSF Grant OCE-1154320 to EBK and KL) and a WHOI Ocean Ventures Fund award to WMJ. The instruments in the WHOI FT-MS Facility were purchased with support from the Gordon & Betty Moore Foundation and NSF. Support for WMJ was provided by a National Defense Science and Engineering Fellowship. Sequencing was performed under the auspices of the US Department of Energy (DOE) JGI Community Science Program (CSP) project (CSP 1685) supported by the Office of Science of US DOE Contract DE-AC02-05CH11231. Additional work related to sample collection and processing was supported by the G. Unger Vetlesen and Ambrose Monell Foundations, the Natural Sciences and Engineering Research Council of Canada (NSERC), the Canadian Institute for Advanced Study (CIFAR) and the Canada Foundation for Innovation through grants awarded to SJH. MPB was supported by a CIFAR Global Scholarship and NSERC postdoctoral fellowship.

